# Vasoinhibin is Generated by the Renin-Angiotensin System

**DOI:** 10.1101/2024.12.04.626908

**Authors:** Francisco Freinet Núñez, Lourdes Siqueiros-Marquez, Elva Adán-Castro, Magdalena Zamora, Juan Pablo Robles, Xarubet Ruiz-Herrera, Thomas Bertsch, Jakob Triebel, Gonzalo Martínez de la Escalera, Carmen Clapp

## Abstract

Vasoinhibin is a fragment of the hormone prolactin (PRL) that inhibits angiogenesis, vasopermeability, and vasodilation. Cathepsin D (CTSD) cleaves the N-teminal of PRL to generate vasoinhibin in the retina of neonate mice as revealed by the CTSD inhibitor, pepstatin A (PA). However, PA also inhibits renin. Because renin is expressed in the retina and the renin-angiotensin system (RAS) gives rise to peptides with positive and negative effects on blood vessel growth and function, we investigated whether renin cleaves PRL to vasoinhibin in the newborn mouse retina and in the circulation. Newborn mouse retinal extracts from wild-type and CTSD-null newborn mice cleaved PRL to a 14 kDa vasoinhibin and such cleavage was prevented by heat-inactivation, PA, and the selective renin inhibitor VTP-27999 suggesting the contribution of renin. In agreement, recombinant renin cleaved different species PRLs to the expected 14 kDa vasoinhibin, a mass consistent with a consensus renin cleavage site located at Leu124-Leu125 in rat and mouse PRLs and at Leu126-Leu127 in human, bovine, and ovine PRLs. Dehydration and rehydration (D/R) in rats increased the levels of renin and PRL in plasma. Further increase in PRL circulating levels by the dopamine D2 receptor blocker, sulpiride, enabled detection of 14 kDa vasoinhibin in D/R rats. Moreover, the incubation of PRL with plasma from D/R rats generated a 14 kDa vasoinhibin that was prevented by VTP-27999. These findings add renin to the list of PRL-cleaving proteases and introduce vasoinhibin as a RAS-mediated mechanism for regulating blood vessel growth and function.

## INTRODUCTION

The proliferation of new blood vessels or angiogenesis underlies the progression of multiple diseases including cancer, rheumatoid arthritis, and vasoproliferative retinopathies making angiogenesis inhibitors promising therapeutics [1,2]. Many inhibitors of angiogenesis are proteolytic fragments of endogenous proteins with no antiangiogenic activity [3,4]. One such inhibitor is vasoinhibin, a proteolytically generated family of fragments of the hormone prolactin (PRL) that inhibits the growth, permeability, and dilation of blood vessels [5,6]. Vasoinhibin isoforms range from 5 to 18 kDa that comprise the first 48 to 159 amino acid residues of PRL depending on the cleavage site of proteases that include thrombin [7], cathepsin D (CTSD) [8], matrix metalloproteases [9], and bone morphogenetic protein 1 [10]. The more studied PRL cleaving protease in tissues is CTSD, an acidic-aspartyl protease that generates vasoinhibin isoforms of 14 to 17 kDa in the secretory granules of anterior pituitary lactotrophs [11], the mammary epithelium [12], the retina [13], and the cartilage [9] of rodents, and in the placenta [14] and pituitary tumors [8] of humans.

Vasoinhibin restricts angiogenesis in the retina and cartilage [9,15], and its levels are modified in peripartum cardiomyopathy [16], preeclampsia [14], rheumatoid arthritis [17], diabetic retinopathy [18], and retinopathy of prematurity (ROP) [19,20]. ROP is a neovascular eye disease that occurs in premature infants having an underdeveloped retinal vascular system [21]. Vasoinhibin is found in the eye of ROP patients [19] and the progression of ROP associates with higher levels of circulating PRL suggestive of lower vasoinhibin levels [20]. Also, the proteolytic conversion of PRL to vasoinhibin decreases in the retina of newborn mice undergoing active vascularization and such conversion is attributed to CTSD, as it is acid pH-dependent and prevented by the CTSD competitive inhibitor, pepstatin A (PA) [13]. However, PA also blocks the activity of renin [22], an aspartyl protease with a catalytic pH range of 5.5 – 7.5 [23]. Renin cleaves circulating angiotensinogen to activate the renin-angiotensin system (RAS), a major regulator of blood pressure, fluid and electrolyte balance, inflammation, and angiogenesis [5,24]. Components of RAS are found in various organs, including the eye [25]. Intra-retinal renin contributes to the local regulation of the vasculature and has been implicated in the pathogenesis of vasoproliferative disorders including ROP [25,26].

Here, we re-evaluated the contribution of CTSD as the only acidic protease generating vasoinhibin in the newborn mouse retina and unveiled renin as a PRL cleaving protease able to generate vasoinhibin that may influence the ocular and systemic vascular properties of RAS.

## METHODS

### Materials

Rat, mouse, bovine, and ovine recombinant PRLs were from the National Hormone and Pituitary Program (AFP4835B, NHPP, National Institute of Diabetes and Digestive and Kidney Diseases [NIDDK] and AF Parlow). Recombinant human PRL generated in *Pichia pastoris* was produced as reported [7]. Antisera against rat PRL (C1), bovine PRL (CL2), and human PRL (HC1) were obtained locally and characterized as reported [19,27]. Monoclonal antibodies (INN-1) that react against the N-terminus of rat PRL were obtained and characterized as reported [15,28]. Recombinant mouse renin generated from the pro-renin activation by trypsin was purchased from R&D Systems (Cat 4277-AS, Minneapolis, MN) and PA was from Sigma-Aldrich (Cat P5318, Saint Louis, MO). VTP-27999 trifluoroacetate (VTP-27999) was from Tocris Bioscience (Cat 5563, Minneapolis, MN).

### Animals

C57BL/6J male and female (1:1) neonate (8 days old) mice wild type (*Ctsd+/+*) or null for CTSD (*Ctsd*-/-) and male Wistar rats (250-300 g) were housed under standard laboratory conditions and cared for in accordance with the US National Research Counciĺs Guide for the Care and Use of Laboratory Animals (Eight Edition, National Academy Press, Washington, D.C., USA). The Bioethics Committee of the Institute of Neurobiology from the National University of Mexico (UNAM) approved all animal experiments. The generation and characterization of C*tsd*-/-mice were previously described [29]. Animals were euthanized by carbon dioxide inhalation and decapitation.

### PRL cleavage analysis

Pools of 5 to 6 neonate mouse retinas were homogenized in 60 µL of lysis buffer (50 mM NaF, 100 mM Na_4_P_2_O_7_, 250 mM sucrose, 0.1 M Tris-HCl, 0.2 M EGTA, 0.2 M EDTA, 1% Igepal, 0.1 M Na_3_VO_4_, pH 7.4) supplemented with a protease inhibitor cocktail (Cat. No. 04693116001, Roche). Recombinant rat PRL (200 ng) in 2 µL of 0.1 mol L^−1^ Tris (pH 7.4) was incubated with 2.5 µg of protein from the retinal homogenate in a final volume of 20 µL incubation buffer at pH 5 (0.1M citrate-phosphate buffer, 0.15 M NaCl) for 24 hours at 37 °C [13]. In some cases, the extract was heat inactivated (95 °C for 30 minutes) or preincubated (30 minutes) with 1.4 µM PA or 10 µM VTP-27999 before adding PRL. In other experiments, recombinant PRLs (200 ng) from different species (mouse, rat, bovine, ovine, and human) were combined with different protein concentrations of pure recombinant mouse renin in a final volume of 20 µL incubation buffer (50 mM NaH_2_PO_4_, 1 M NaCl) adjusted to pH 5.5 or 7.5 and incubated for 24 hours at 37 °C. In some cases, the renin was heat-inactivated or preincubated with PA before adding the PRLs. Reactions were stopped by the addition of reducing Laemmli sample buffer and 15% SDS-PAGE fractionation or by storing the samples at -70 °C.

### Western blot

The products of PRL cleavage resolved under reducing 15% SDS-PAGE were transferred to nitrocellulose membranes and probed overnight with a 1:500 dilution of antisera against rat PRL (C-1), human PRL (HC1), bovine PRL (CL-2) or of monoclonal antibodies (INN-1) that react with the N-terminus of rat PRL. Detection used anti-rabbit or anti-mouse secondary antibodies linked to alkaline phosphatase (Jakson ImmunoResearch) and the alkaline phosphatase conjugate substrate kit (Bio-Rad). The ImageJ sofware analysis program (NIH) evaluated optical density values.

### Alignment of vertebrate PRL sequences

The amino acid sequences of mammalian PRLs (human, simian, bovine, ovine, rat, mouse) and avian (*Gallus gallus*), reptilian (*Pelodiscus sinensis*), amphibian (*Xenopus laevis*), and pisces (*Danio rerio*) PRLs were obtained from GenBank. The alignment of these sequences was carried out using the MUSCLE program (Multiple Sequence Comparison by Log-Expectation) of the European Molecular Biology Laboratory (EMBL) – European Bioinformatics Institute (EBI). The alignment focused particularly on the region comprised by the sequence NKRLLEGM of the PRL molecule. The conservation percentage of this sequence was obtained using Jalview [30] and the sequence logo was designed using WebLogo [31].

### Real time RT-PCR

Total RNA was isolated from neonate mouse retinas and from rat’s kidneys using Qiagen RNeasy Mini Kit (Qiagen, Germantown, MD) and reverse transcribed with the High-Capacity cDNA Reverse Transcription Kit (Applied Biosystems, Foster City, CA). Maxima SYBR Green/ROX qPCR Master Mix (Thermo Scientific, Waltham, MA) quantified PCR products in a 10 μL final reaction volume containing template (25 ng) and 0.5 µmolL^−1^ of each primer pair for mouse *Ctsd*: 5’-AAGCTGGTGGACAAGAACAT-3’ and 5’-TTTCGAGTGACGTTCAGGTA-3’; mouse *Renin*: 5’-GTTCATCCTTTATCTCGGCT-3’ and 5’-AATCCAATGCGATTGTTATG -3’; rat *Renin*: 5’-ATGGGCGGGAGGAGGATGCC-3’ and 5’-TTAGCGGGCCAAGGCGAACCC-3’; and hypoxanthine guanine phosphoribosyl transferase (*Hprt*): 5’-TTGCTGACCTGCTGGATTAC-3’ and 5’-GTTGAGAGATCATCTCCACC-3’. Amplification was for 10 seconds at 95 °C, 30 seconds at each primer pair-specific annealing temperature, and 30 seconds at 72 °C for 40 cycles. The mRNA expression levels were calculated using the 2^−ΔΔCT^ method and normalized to the housekeeping gene *Hprt*.

### Dehydration / rehydration in rats

Wistar rats were subjected or not (control) to a previously reported dehydration and rehydration (D/R) protocol [32] in which rats were deprived of water for 48 hours, returned to water for 3 hours, and euthanized immediately after. Kidneys and plasmas were collected and stored at -70 °C to evaluate the renal expression of renin and the circulating levels of renin and PRL. To favor detection of circulating vasoinhibin, the systemic levels of PRL were further elevated in control or D/R rats by the intraperitoneal injection of sulpiride (20 mg/kg, Dogmatil, Sanofi) [33] 6 hours before sacrifice (i.e., 3 hours before regaining access to water).

### ELISAs

The Rat Renin 1 ELISA Kit (RayBiotech Life, Norcross, GA) and the previously reported ELISA method [34] measured renin and PRL levels in plasma, respectively.

### Immunoprecipitation – Western blot

The immunoprecipitation-Western blot method analyzed the circulating levels of vasoinhibin and was carried out as reported [7]. Briefly, pools of plasma samples (500 µL) from three rats per each group were incubated overnight at 4 °C with 2 µL of C-1 anti-rat PRL antiserum, followed by a 2-hour incubation with protein-A Sepharose beads (40 µL, Sigma-Aldrich). The samples were centrifuged and washed three times with PBS (pH 7.4). The final pellet was resuspended in reducing Laemmli buffer, heated at 97 °C for 15 minutes under agitation, centrifuged for 5 minutes, and the supernatant subjected to 15% SDS/PAGE-Western blot probed with the C-1 anti-rat PRL antiserum.

### PRL cleavage with plasma

Human PRL was added to a final concentration of 4 µM in 120 µL of plasma from D/R rats with or without 10 µM VTP-27999, incubated for 24 hours at 37 °C. One hundred µL of the reaction mixture were incubated overnight with 2 µL of HC1 anti-human PRL antiserum followed by a 2-hour incubation with 40 µL protein-A Sepharose beads. Samples were then centrifuged and the immunoprecipitate subjected to 15% SDS-PAGE-Western blots probed with the HC1 anti-human PRL antiserum. The incubation of human PRL with plasma from D/R rats allowed assaying the cleaving activity of circulating renin without the confounding intervention of endogenous rat hormones (i.e., rat PRL and vasoinhibin which do not react with anti-human PRL antibodies).

### Statistical analysis

Statistical analysis was performed using GraphPad Prism version 8.01 for Windows, GraphPad Sofware (Boston, MA, www.graphpad.com). The values are expressed as mean ± SEM. The unpaired two-tailed Student’s *t*-test evaluated differences between two groups. The threshold for significance was set at P ≤ 0.05.

## RESULTS

### CTSD is not the only acidic protease generating vasoinhibin in the retina of newborn mice

To re-evaluate CTSD as the only responsible acidic protease cleaving PRL to vasoinhibin in the retina of newborn mice, rat PRL was incubated with an acidic (pH 5.0) retinal extract from *Ctsd*+/+ or *Ctsd*-/- mice at postpartum day 8 using the same proportions shown to be effective for PRL cleavage in the wild type condition [13]. The cleaved PRL products were evaluated by Western blot. A representative Western blot probed with an anti-rat PRL antiserum (C1) showed that rat PRL incubated in the absence of retinal extract, migrates as a major 23 kDa band (the molecular mass of intact PRL) and a minor ∼13 kDa band (Figure 1A, lane 1). The 13 kDa PRL isoform was present in the recombinant rat PRL standard before incubation and only in the batch used for the experiments in Figures 1A and B. The cleavage likely occurred during the generation and/or purification of the recombinant hormone and, thereby, the 13 kDa isoform was considered irrelevant for the present study.

**Figure 1.**
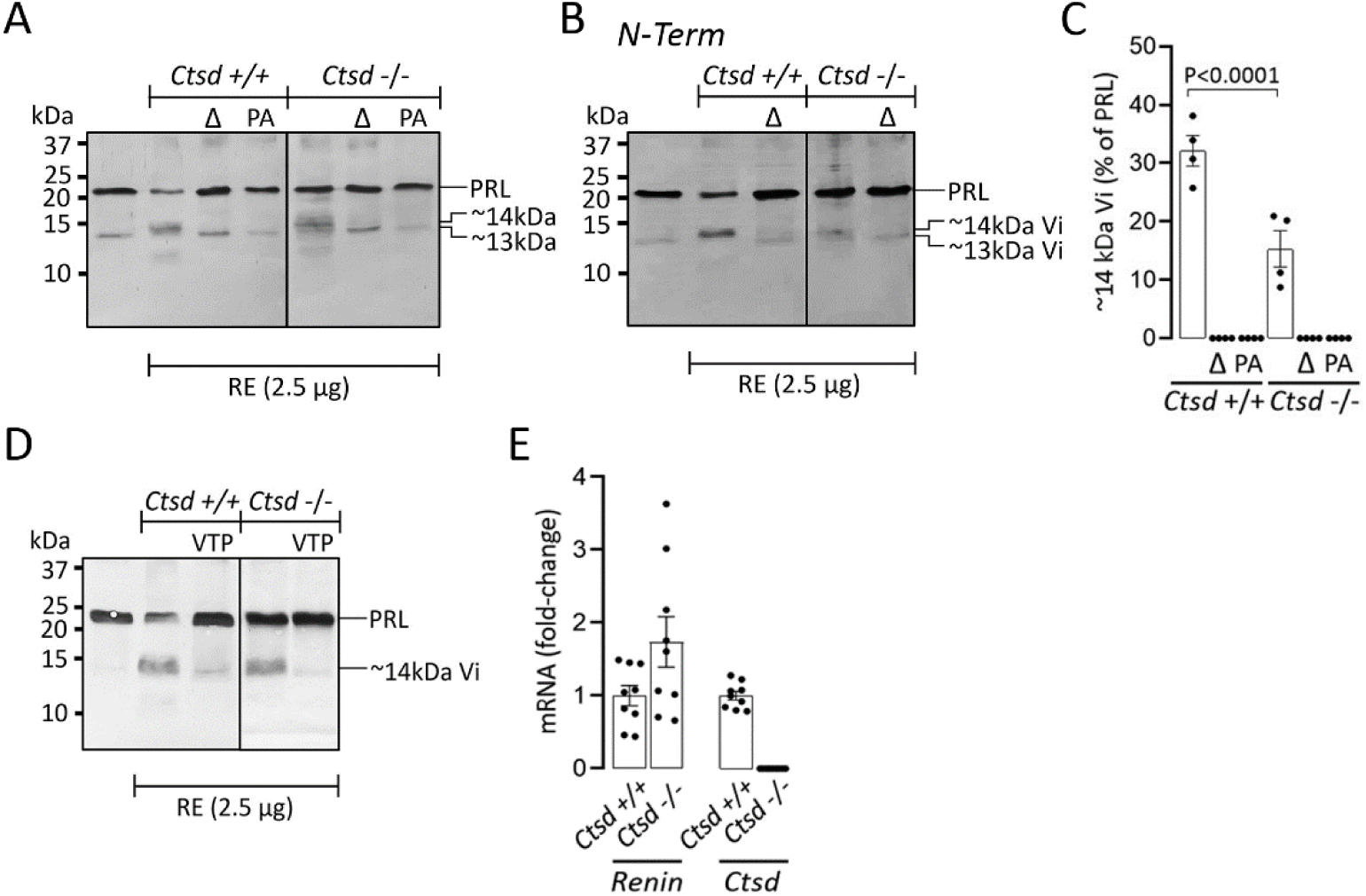
Cathepsin D (CTSD) is not the only acidic protease cleaving PRL to vasoinhibin in the retina of newborn mice. **(A)** Representative Western blot probed with an anti-rat PRL antiserum showing the cleaved products derived from the incubation of rat PRL without or with 2.5 µg of protein from newborn mouse retinal extracts (RE) of *Ctsd* +/+ or *Ctsd* -/- mice carried out in the absence or presence of heat-inactivated RE (Δ) or pepstatin A (PA). PRL and PRL fragments of ∼14 and ∼13 kDa are indicated. The numbers on left indicate the weight (kDa) of molecular markers. **(B)** Representative Western blot, like in **A**, but probed with monoclonal antibodies that react with the N-terminus of rat PRL (N-Term). PRL and vasoinhibins (Vi) of ∼14 and ∼13 kDa are indicated. The numbers on left indicate the weight (kDa) of molecular markers. **(C)** Densitometric values of the ∼14 kDa Vi band plotted relative to the values of the PRL band incubated without retinal extract. Values are means ± SEM from 3 independent experiments. **(D)** Representative Western blot probed with an anti-rat PRL antiserum showing the cleaved products derived from the incubation of rat PRL without or with 2.5 µg of protein from RE of newborn *Ctsd*+/+ or *Ctsd*-/- mice in the absence or presence of VTP-27999 (VTP). PRL and ∼14 kDa vasoinhibin (Vi) are indicated. The numbers on left indicate the weight (kDa) of molecular markers. **(E)** Expression of mRNA by RT-PCR of *Renin* and *Ctsd* in retina of newborn *Ctsd*+/+ or *Ctsd*-/- mice calculated relative to the control gene *Hprt*. Values are means ± SEM (n=9).

As previously reported [13], incubation with retinal extracts from wild type newborn mice, resulted in the partial conversion of PRL to a main ∼14 kDa vasoinhibin-like protein (Figure 1A, lane 2) and such conversion was abolished by the heat inactivation of the extract or the addition of PA (Figure 1A, lanes 3 and 4). The incubation of PRL with retinal extracts from *Ctsd*-/- mice did not eliminate the generation of the ∼14 kDa vasoinhibin-like protein (Figure 1A, lane 5), whereas heat-inactivation or PA did (lanes 6 and 7). Because the functional determinant defining vasoinhibin is located within the first 48 amino acid residues of PRL [35], Western blots probed with monoclonal antibodies directed against the N-terminus of PRL verified the vasoinhibin nature of the ∼14 kDa vasoinhibin-like protein (Figure 1B). Optical density values from independent experiments, evaluated with N-terminal anti-PRL antibodies and expressed relative to those of the PRL incubated in the absence of retinal extract, revealed that ∼50% of the ∼14 kDa vasoinhibin was generated in the absence of CTSD (Figure 1C). These findings indicate that CTSD is not the only acidic protease generating vasoinhibin in the newborn mouse retina.

### Renin contributes to the generation of vasoinhibin in the retina of newborn mice

Because PA blocks the activity of renin [22] and renin is active at acid pH [23], we next investigated whether renin cleaves PRL to vasoinhibin in the retina by testing the effect of VTP-27999, a highly selective and direct renin inhibitor [36] (Figure 1D). VTP-27999 nearly abolished the cleavage of PRL to the ∼14 kDa vasoinhibin by the retinal extracts from both *Ctsd*+/+ and *Ctsd*-/- newborn mice (Figure 1D). Such inhibition argues in favor of renin playing a role in the generation of vasoinhibin in the neonate mouse retina, in as much as at the concentration used (10 µM), VTP-27999 is a weak inhibitor of other aspartyl proteases, including CTSD [36]. Furthermore, we evaluated the mRNA expression of *Renin* in retinas from *Ctsd*+/+ and *Ctsd*-/- mice at postpartum day 8. *Renin* and *Ctsd* were expressed in the newborn mouse retinas under the wild type condition and, as expected, only *Renin* was expressed in mice null for CTSD (Figure 1E).

### Purified renin cleaves PRL into a ∼14 kDa vasoinhibin

Western blots probed with the anti-rat PRL antiserum showed that recombinant mouse renin cleaves rat PRL in a dose-dependent manner to a ∼14 kDa fragment both at pH 5.5 and pH 7.5 (Figure 2A), the catalytic range of renin activity [23]. Optical density values of the ∼14 kDa fragment, relative to those of PRL in the absence of renin, indicated that the cleavage of PRL was equally effective at acid and neutral pHs (Figure 2B). Specificity was verified by the lack of PRL cleavage after the heat inactivation of renin or the addition of PA (Figure 2C). The vasoinhibin identity of the ∼14 kDa PRL fragment was verified by its reaction with the N-terminal anti-PRL monoclonal antibodies (Figure 2D). These findings showed that renin cleaves PRL into a ∼14 kDa vasoinhibin and, thereby, support its contributing role as a PRL cleaving protease generating vasoinhibin in the newborn mouse retina.

**Figure 2.**
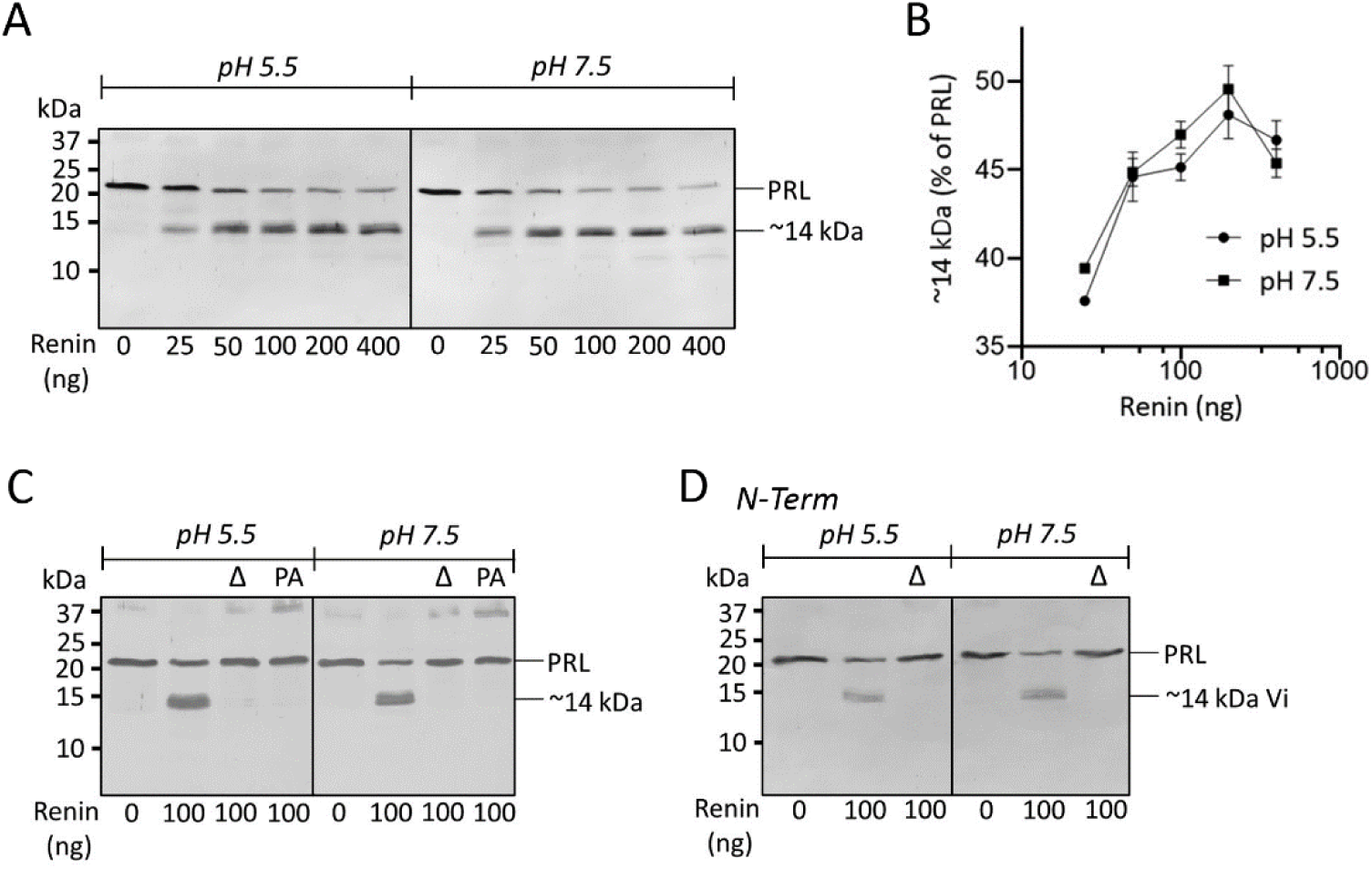
Renin cleaves PRL to generate a ∼14 kDa vasoinhibin at acid and neutral pH. **(A)** Representative Western blots probed with an anti-rat PRL antiserum showing the PRL cleaved products generated after incubation of rat PRL in the absence (0) or presence of increasing amounts of recombinant mouse renin at pH 5.5 or pH 7.5. PRL and a PRL fragment of ∼14 kDa are indicated. **(B)** Densitometric values of the ∼14 kDa PRL band relative to the values of the PRL band incubated without renin at pH 5.5 and pH 7.5. Values are means ± SEM. **(C)** Representative Western blot probed with an antirat PRL antiserum of the cleaved products derived from the incubation of rat PRL with renin heat-inactivated (Δ) or not, or with renin and pepstatin A (PA) at pH 5.5 or pH 7.5. PRL and a PRL fragment of ∼14 kDa are indicated. (**D**) Representative Western blot, like **C**, but probed with monoclonal antibodies that react with the N-terminus of rat PRL (N-Term). PRL and the ∼14 kDa vasoinhibin (Vi) are indicated. The numbers on left indicate the weight (kDa) of molecular markers.

### PRL cleavage by renin is evolutionarily conserved

The relevance of renin as a PRL protease is supported by its ability to cleave PRLs from different mammalian species. Renin cleaved rat, mouse, ovine, bovine, and human PRL into the ∼14 kDa vasoinhibin (Figure 3A), implying a renin cleavage site within a conserved region of PRL. Except for the renin cleavage site in human angiotensinogen (Leu10–Val11), renin cleaves angiotensinogen of most mammalian and non-mammalian vertebrates between two Leu residues (Leu10–Leu11) to generate the decapeptide, angiotensin I [37] (Figure 3B). Rat and mouse PRLs have the renin consensus cleavage site at Leu124–Leu125, whereas bovine / ovine and human PRLs at Leu126–Leu127. Sequences comprising residues 1–124 and 1–126 have a calculated mass of 14 and 14.3 kDa, respectively, the estimated molecular mass of the vasoinhibin generated by renin from the above mammalian PRLs (Figure 3A) and in the neonate mouse retina (Figure 1). The renin proposed cleavage sites are within the NKRLLEG sequence, a highly conserved region of the PRL molecule, where even in non-mammalian vertebrates is nearly identical (Figure 3B). Multiple alignment of vertebrate PRLs revealed an almost absolute conservation of the sequences on the C-terminal side of the Leu–Leu bond in mammalian PRLs and in both the N-terminal and C-terminal sides in non-mammalian PRLs (Figure 3C). These observations suggest that the generation of vasoinhibin by renin is conserved throughout evolution.

**Figure 3.**
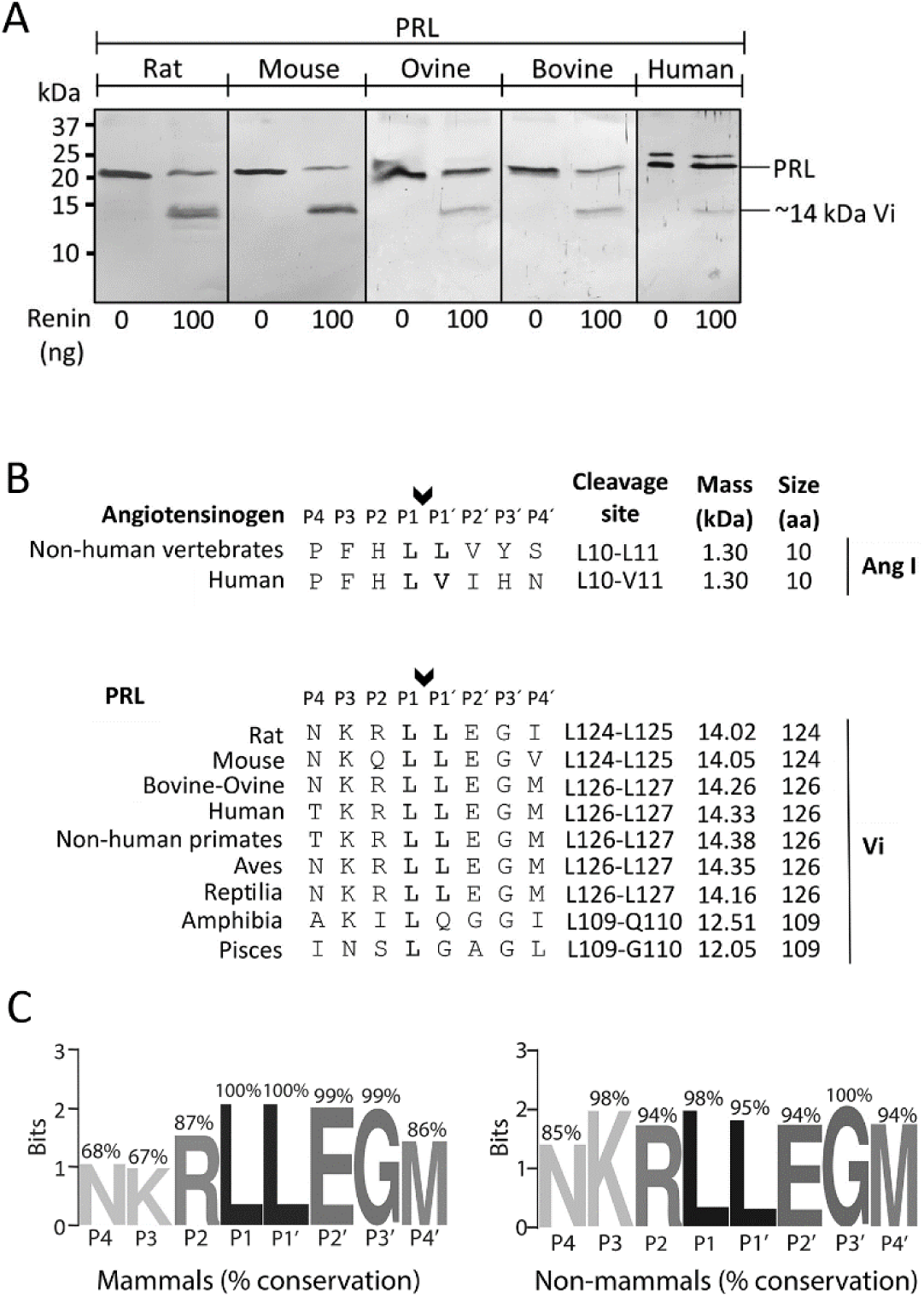
Renin cleaves PRL to generate a ∼14 kDa vasoinhibin at a consensus cleavage site conserved throughout evolution. **(A)** Representative Western blots probed with antisera against rat, mouse, ovine, and human PRLs showing the cleaved products generated after the incubation of the different mammalian PRLs with recombinant mouse renin at pH 7.5. PRL and the ∼14 kDa vasoinhibin (Vi) are indicated. The numbers on left indicate the weight (kDa) of molecular markers. **(B)** Alignment of the renin natural substrate, angiotensinogen, at the renin consensus cleavage site (L10−L11 in non-human vertebrates and L10−V11 in humans) that generates angiotensin I (Ang I). Alignment of the N-terminal region from mammalian and non-mammalian PRLs at the nearly absolute conserved NKRLLEGI sequence that contains consensus cleavage sites (L124−L125 in rodents and L126−L127 in most other vertebrates) for renin that upon cleavage generate vasoinhibins (Vi) with a calculated mass of ∼14 kDa. Amino acid residues in substrates are numbered outward from the cleavage site occurring between P1 and P1’ (arrow). **(C)** Representations of the amino acid frequencies and level of conservation in the P4 − P4’ cleavage sequence of renin in mammalian and non-mammalian PRLs.

### Endogenous renin generates vasoinhibin in the circulation

To further analyze the physiological relevance of the PRL cleavage by renin, we tested whether the upregulation of renin in response to D/R [32] could result in the generation of circulating vasoinhibin. First, we showed that rats dehydrated for 48 hours and returned to water for 3 hours upregulated *Renin* mRNA expression in the kidney (Figure 4A) and confirmed the previously observed [32] higher levels of circulating renin using the same protocol (Figure 4B). We also found that D/R increases by 4-fold the levels of PRL in plasma (20.4 ± 6.1 ng/mL vs. 5.1 ± 1.7 ng/mL) (Figure 4C). In the absence of a quantitative assay for vasoinhibin, its circulating levels can only be determined semi-quantitatively by immunoprecipitation followed by Western blot. However, this method is not sensitive enough to detect circulating vasoinhibin unless higher PRL levels are present, like under sulpiride-induced hyperprolactinemia [33], under the notion that increasing the substrate (PRL) results in more product (vasoinhibin) and, thereby, favors vasoinhibin detection. Sulpiride increased to a similar level the circulating values of PRL determined by ELISA in both, control (143.9 ± 12.3 ng/mL) and D/R rats (171.2 ± 22.6 ng/mL) (Figure 4D). Figure 4E shows the immunoprecipitation-Western blot analysis of plasma samples representative of three independent experiments. In the absence of sulpiride, the plasma from control and D/R rats contained PRL but vasoinhibin-like proteins were not detected. In agreement with the ELISA, sulpiride elevated PRL to a similarly high level in control and D/R rats, however only the plasma from D/R contained a ∼14 kDa vasoinhibin like protein. To evaluate circulating renin as the responsible protease generating 14 kDa vasoinhibin in the plasma of D/R rats, human PRL was incubated without or with the plasma of D/R rats in the presence and absence of VTP-27999 (Figure 4F). Representative Western blot showing the immunoprecipitation-Western blot analysis of human PRL incubated without plasma (lane 1) and the plasma from D/R rats incubated without human PRL (lane 2) or with human PRL in the absence (lane 3) or presence (lane 4) of VTP-27999. The plasma from D/R rats generated a 14 kDa immunoreactive human PRL (14 kDa vasoinhibin) and such generation was prevented by the specific inhibition of renin with VTP-27999 (Figure 4F).

**Figure 4.**
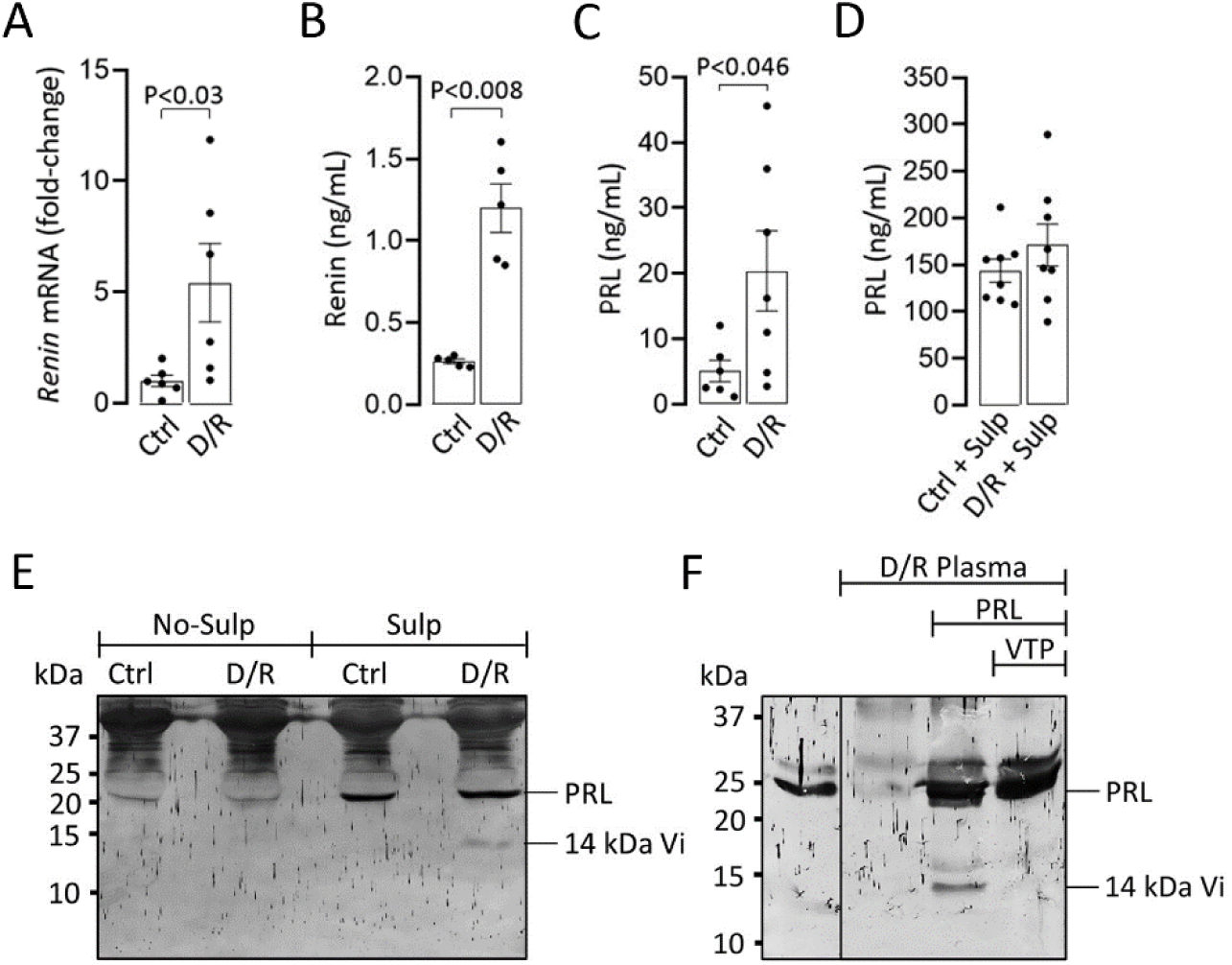
Renin generates vasoinhibin in the circulation. Renal expression of *Renin* mRNA (**A**) and plasma levels of renin (**B**) and PRL (**C**) in control (Ctrl) rats and in rats subjected to dehydration/rehydration (D/R). PRL levels were also determined in Ctrl and D/R rats treated with sulpiride (Sulp) (**D**). Values are means ± SEM. (**E**) Representative immunoprecipitation -Western blot evaluating PRL and vasoinhibin (Vi) levels in plasma samples from Ctrl and D/R rats treated or not with sulpiride. PRL and a ∼14 kDa Vi are indicated. The numbers on left indicate the weight (kDa) of molecular markers. (**F**) Representative immunoprecipitation-Western blot showing the generation of 14 kDa vasoinhibin (Vi) following the incubation of human PRL without or with the plasma from D/R rats in the absence or presence of VTP-27999 (VTP). PRL and Vi are indicated. Numbers on left indicate the weight (kDa) of molecular markers.

These findings suggest that renin released by changes in blood volume due to D/R cleaved circulating PRL to 14 kDa vasoinhibin and are proof of concept that renin generates vasoinhibin in association with the systemic activation of RAS.

## DISCUSSION

Proteolytic processing is a mechanism for creating functional diversity in peptide hormones [5]. PRL acquires inhibitory properties on blood vessels after undergoing proteolytic cleavage to vasoinhibin, a family of PRL fragments that inhibits angiogenesis, vascular permeability, and vasodilation [5,6]. The cleavage of angiotensinogen by renin initiates the proteolytic cascade that generates RAS peptides controlling blood pressure, body fluid homeostasis, inflammation, and angiogenesis [5,24,38]. Here, we introduce renin as a PRL cleaving protease that can act locally and systemically to produce vasoinhibin able to contribute to the vascular actions of RAS. The generation of vasoinhibin depends on the levels of PRL and the activity of PRL cleaving proteases regulated at the hypothalamus, the pituitary, and the target tissues, defining the PRL/vasoinhibin axis [39]. This axis helps restrict angiogenesis in ocular and joint tissues [9,15] and is disrupted in vasoproliferative retinopathies [18,20], rheumatoid arthritis [17], peripartum cardiomyopathy [16], and preeclampsia [14]. The PRL/vasoinhibin axis was recently studied in ROP, a potentially blinding neovascular eye disease that occurs in premature babies [21]. The circulating levels of PRL were very high before and during ROP and were not followed by a concomitant increase in vasoinhibin [20] to suggest reduced vasoinhibin generation. In agreement, the cleavage of PRL to a ∼14 kDa vasoinhibin-like protein was lower in newborn mouse retinas undergoing active neovascularization and was attributed to CTSD because it depended on acidic pH and was blocked by PA [13].

Here, we have challenged CTSD as the only PRL cleaving acidic protease in the newborn mouse retina. We identified the ∼14 kDa vasoinhibin-like protein as vasoinhibin by showing that it is an N-terminal fragment of PRL containing the functional anti-angiogenic motif [35]. Furthermore, we found that 50% of the ∼14 kDa vasoinhibin was generated in retinas from mice null for CTSD. In search of a contributing enzyme, we unveiled renin, another aspartyl protease active at acid pH (and neutral pH) that is inhibited by PA [22] and expressed in the retina of rodents and humans [26,40,41]. We showed that the highly selective renin inhibitor VTP-27999 [36] nearly abolished the generation of 14 kDa vasoinhibin following PRL incubation with retinal extracts from newborn mice null or not for CTSD, that both *Ctsd*+/+ and *Ctsd*-/- newborn mouse retinas express *Renin*, and that recombinant renin cleaves PRL to the expected 14 kDa vasoinhibin.

Uncovering PRL as substrate for renin is remarkable since the only known substrate for renin is angiotensinogen. Renin cleaves the Leu–Leu bond in position 10–11 of angiotensinogens, from most species of vertebrates, to generate the decapeptide angiogensin I [37]. PRL of mammalian and non-mammalian vertebrates has a Leu–Leu bond, within the nearly absolute conserved sequence NKRLLEGM, that when cleaved generates a 14 kDa vasoinhibin (residues 1–124 in rodents and 1–126 in most other vertebrates) and renin cleaved different mammalian PRLs into the 14 kDa vasoinhibin. The PRL/vasoinhibin axis exerts a wide diversity of actions on the development, growth, and reproduction of vertebrates that are fueled by its effects on blood vessels [42]. The evolutionary conservation of the putative renin cleavage site suggests that renin produces vasoinhibin in all vertebrates. The conservation of angiotensin peptides throughout evolution [43] prompted investigating whether the generation of vasoinhibin by renin could be linked to the activation of RAS.

The PRL/vasoinhibin axis and RAS can be functionally interconnected by renin. A D/R protocol, that upregulates the levels of active renin and angiotensin II in the circulation [32], increased the systemic levels of renin, PRL, and vasoinhibin. The upregulation of systemic PRL following water deprivation was first reported nearly 5 decades ago [44] and it can occur under apparent stress-free conditions [45]. Higher PRL levels promote the conversion to vasoinhibin by providing more substrate for cleavage.

Hyperprolactinemic mice overexpressing PRL in the liver have enhanced levels of circulating vasoinhibin [46] and pharmacologically induced hyperprolactinemia results in higher levels of vasoinhibin in ocular tissues and fluids of rats [33] and humans [47].

However, the increase in systemic PRL induced by D/R was not enough to detect the systemic conversion to vasoinhibin by renin. Therefore, we further elevated circulating PRL levels using sulpiride, a dopamine D2 receptor blocker that hinders the hypothalamic dopaminergic inhibition of PRL secretion by the anterior pituitary gland [48]. Plasma samples from sulpiride-treated D/R rats, but not from sulpiride-treated controls, contained a 14 kDa vasoinhibin-like protein, like the vasoinhibin produced by pure renin. Although other known or unknown PRL cleaving proteases cannot be ruled out, renin is the likely protease responsible for the generation of vasoinhibin in the circulation of hyperprolactinemic rats subjected to D/R. D/R upregulates systemic renin and hyperprolactinemia did not result in higher vasoinhibin levels in the absence of D/R.

Also, there is no clear evidence that matrix metalloproteases and/or CTSD increase in the circulation following D/R and CTSD is acid pH-dependent and has no activity at the neutral pH found in the circulation. Furthermore, performing the study in *Ctsd*-/- mice to eliminate a putative CTSD contribution is not feasible as these animals die young (at 26 days of age) due to progressive atrophy of the intestinal mucosa and destruction of lymphoid cells [29]. The contribution of circulating renin to the generation of vasoinhibin was decisively demonstrated by showing that human PRL incubated with the plasma from D/R rats was partially converted to the 14 kDa vasoinhibin and that such conversion was prevented by the specific inhibition of renin with VTP-27999.

Altogether, the present findings support that, like in RAS, renin released by changes in blood volume activates the PRL/vasoinhibin axis. A major question is whether the PRL/vasoinhibin axis contributes to RAS functions. RAS acts systemically to regulate blood pressure and water balance, and operates locally, within a tissue / organ, to control blood flow, inflammation, and angiogenesis [38]. PRL and vasoinhibin share functional properties with RAS. The PRL/vasoinhibin axis is upregulated in hypertensive patients [14,46] and elevates blood pressure [46], reduces blood flow [49], and inhibits the relaxation of coronary vessels and aortic segments [50]. These effects are brought about by the vasoinhibin mediated impairment of endothelial nitric oxide synthase (eNOS) NO production [50]. Because the pressure and volume of blood are closely interrelated, it is not surprising that the PRL/vasoinhibin axis is upregulated by water deprivation and uptake (present data) [44]. Also, hyperprolactinemia potentiates the dipsogenic effect of angiotensin II [45] and, like angiotensin II [51], both PRL and vasoinhibin stimulate the release of vasopressin from the hypothalamo-neurohyphyseal system [52]. With respect to local actions, vasoinhibin acts on endothelial cells to inhibit the signaling pathways (Ras-Raf-MAPK, Ras-Tiam1-Rac1-Pac1, PI3K-Akt, and PLCγ-IP3-eNOS) activated by several proangiogenic, vascular permeability, and vasodilation factors (VEGF, bFGF, bradykinin, interleukin 1β, acetylcholine) [5,6]. Likewise, vasoinhibin activates, by itself, the NF-kB pathway in endothelial cells [53] and fibroblasts [54] to trigger the expression of adhesion molecules and inflammatory mediators resulting in leukocyte infiltration and inflammation. The functional interconnection between the PRL/vasoinhibin axis and RAS is supported by their involvement in the pathophysiology of vasoproliferative retinopathies, including ROP [19,20,25,26]. The components of both systems are found in the retina, are disrupted under retinal vascular alterations, and angiotensin peptides, PRL, and vasoinhibin can worsen or improve vasoproliferative retinopathies [5,18,20,25,26,55].

In conclusion, the present study unveils renin as a PRL cleaving protease able to act locally and systemically to produce vasoinhibin under conditions linked to the activation of RAS. The functional similarities between the PRL/vasoinhibin axis and RAS highlight the need to address the concomitant generation by renin of vasoinhibin and angiotensin peptides with the final goal of understanding their contribution to vascular diseases and the development of new treatments.

## ABBREVIATIONS

PRL: Prolactin
CTSD: Cathepsin D
PA: Pepstatin A
RAS: Renin-Angiotensin System
D/R: dehydration/rehydration.

## ACKNOWLEDGEMENTS

We thank Paul Saftig for providing the CTSD null mice and Jose Fernando García-Rodrigo, Fernando Macías, Fernando López Barrera, Alejandra Castilla, Martín García, and María Antonieta Carbajo for their excellent technical assistance.

## AUTHORSHIP CONTRIBUTION STATEMENT

F. F. N. conceived and designed experiments, acquired and analyzed data, helped draft the manuscript; L. S.-M. designed experiments, acquired and analyzed data, and supervised the study; E. A.-C acquired and analyzed data; M. Z. designed experiments, acquired and analyzed data; J. P. R. and X. R.-H. acquired and analyzed data; T. B, and J. T provided resources; G. M. de la E., supervised and analyzed the data and secured funding; C. C., conceived, designed, supervised the study, analyzed the data, secured funding, and wrote the manuscript. All authors critically reviewed and approved the submitted version of the manuscript.

## REFERENCES

[1] P. Carmeliet, Angiogenesis in health and disease, Nat Med 9 (2003) 653–660. 10.1038/nm0603-653.

[2] A. Fallah, A. Sadeghinia, H. Kahroba, A. Samadi, H.R. Heidari, B. Bradaran, S. Zeinali, O. Molavi, Therapeutic targeting of angiogenesis molecular pathways in angiogenesis-dependent diseases, Biomedicine & Pharmacotherapy 110 (2019) 775–785. 10.1016/j.biopha.2018.12.022.

[3] P. Nyberg, L. Xie, R. Kalluri, Endogenous Inhibitors of Angiogenesis, Cancer Research 65 (2005) 3967–3979. 10.1158/0008-5472.CAN-04-2427.

[4] Y. Cao, Endogenous angiogenesis inhibitors and their therapeutic implications, The International Journal of Biochemistry & Cell Biology 33 (2001) 357–369. 10.1016/S1357-2725(01)00023-1.

[5] C. Clapp, S. Thebault, M.C. Jeziorski, G. Martínez De La Escalera, Peptide Hormone Regulation of Angiogenesis, Physiological Reviews 89 (2009) 1177–1215. 10.1152/physrev.00024.2009.

[6] C. Clapp, S. Thebault, Y. Macotela, B. Moreno-Carranza, J. Triebel, G. Martínez de la Escalera, Regulation of Blood Vessels by Prolactin and Vasoinhibins, in: P. Diakonova Maria (Ed.), Recent Advances in Prolactin Research, Springer International Publishing, Cham, 2015: pp. 83–95. 10.1007/978-3-319-12114-7_4.

[7] M. Zamora, J.P. Robles, M.B. Aguilar, S. de J. Romero-Gómez, T. Bertsch, G. Martínez de la Escalera, J. Triebel, C. Clapp, Thrombin Cleaves Prolactin Into a Potent 5.6-kDa Vasoinhibin: Implication for Tissue Repair, Endocrinology 162 (2021) bqab177. 10.1210/endocr/bqab177.

[8] D. Piwnica, P. Touraine, I. Struman, S. Tabruyn, G. Bolbach, C. Clapp, J.A. Martial, P.A. Kelly, V. Goffin, Cathepsin D Processes Human Prolactin into Multiple 16K-Like N-Terminal Fragments: Study of Their Antiangiogenic Properties and Physiological Relevance, Molecular Endocrinology 18 (2004) 2522–2542. 10.1210/me.2004-0200.

[9] Y. Macotela, M.B. Aguilar, J. Guzmán-Morales, J.C. Rivera, C. Zermeño, F. López-Barrera, G. Nava, C. Lavalle, G.M. de la Escalera, C. Clapp, Matrix metalloproteases from chondrocytes generate an antiangiogenic 16 kDa prolactin, Journal of Cell Science 119 (2006) 1790–1800. 10.1242/jcs.02887.

[10] G. Ge, C.A. Fernández, M.A. Moses, D.S. Greenspan, Bone morphogenetic protein 1 processes prolactin to a 17-kDa antiangiogenic factor, Proc. Natl. Acad. Sci. 104 (2007) 10010–10015. 10.1073/pnas.0704179104.

[11] M.E. Cruz-Soto, G. Cosío, M.C. Jeziorski, V. Vargas-Barroso, M.B. Aguilar, A. Cárabez, P. Berger, P. Saftig, E. Arnold, S. Thebault, G. Martínez de la Escalera, C. Clapp, Cathepsin D Is the Primary Protease for the Generation of Adenohypophyseal Vasoinhibins: Cleavage Occurs within the Prolactin Secretory Granules, Endocrinology 150 (2009) 5446–5454. 10.1210/en.2009-0390.

[12] M. Lkhider, R. Castino, E. Bouguyon, C. Isidoro, M. Ollivier-Bousquet, Cathepsin D released by lactating rat mammary epithelial cells is involved in prolactin cleavage under physiological conditions, Journal of Cell Science 117 (2004) 5155–5164. 10.1242/jcs.01396.

[13] M. Vázquez-Membrillo, L. Siqueiros-Márquez, F.F. Núñez, N. Díaz-Lezama, E. Adán-Castro, G. Ramírez-Hernández, N. Adán, Y. Macotela, G. Martínez de la Escalera, C. Clapp, Prolactin stimulates the vascularisation of the retina in newborn mice under hyperoxia conditions, J. Neuroendocrinol. 32 (2020) e12858. 10.1111/jne.12858.

[14] C. González, A. Parra, J. Ramírez-Peredo, C. García, J.C. Rivera, Y. Macotela, J. Aranda, M. Lemini, J. Arias, F. Ibargüengoitia, G.M. de la Escalera, C. Clapp, Elevated vasoinhibins may contribute to endothelial cell dysfunction and low birth weight in preeclampsia, Lab. Invest. 87 (2007) 1009–1017. 10.1038/labinvest.3700662.

[15] J. Aranda, J.C. Rivera, M.C. Jeziorski, J. Riesgo-Escovar, G. Nava, F. López-Barrera, H. Quiróz-Mercado, P. Berger, G. Martínez de la Escalera, C. Clapp, Prolactins Are Natural Inhibitors of Angiogenesis in the Retina, Invest. Ophthalmol. Vis. Sci. 46 (2005) 2947–2953. 10.1167/iovs.05-0173.

[16] D. Hilfiker-Kleiner, K. Kaminski, E. Podewski, T. Bonda, A. Schaefer, K. Sliwa, O. Forster, A. Quint, U. Landmesser, C. Doerries, M. Luchtefeld, V. Poli, M.D. Schneider, J.-L. Balligand, F. Desjardins, A. Ansari, I. Struman, N.Q.N. Nguyen, N.H. Zschemisch, G. Klein, G. Heusch, R. Schulz, A. Hilfiker, H. Drexler, A Cathepsin D-Cleaved 16 kDa Form of Prolactin Mediates Postpartum Cardiomyopathy, Cell 128 (2007) 589–600. 10.1016/j.cell.2006.12.036.

[17] C. Clapp, N. Adán, M.G. Ledesma-Colunga, M. Solís-Gutiérrez, J. Triebel, G. Martínez de la Escalera, The role of the prolactin/vasoinhibin axis in rheumatoid arthritis: an integrative overview, Cell. Mol. Life Sci. 73 (2016) 2929–2948. 10.1007/s00018-016-2187-0.

[18] J. Triebel, T. Bertsch, C. Clapp, Prolactin and vasoinhibin are endogenous players in diabetic retinopathy revisited, Front. Endocrinol. 13 (2022) 994898. 10.3389/fendo.2022.994898.

[19] Z. Dueñas, J.C. Rivera, H. Quiróz-Mercado, J. Aranda, Y. Macotela, P.M. de Oca, F. López-Barrera, G. Nava, J.L. Guerrero, A. Suarez, M. De Regil, G.M. de la Escalera, C. Clapp, Prolactin in Eyes of Patients with Retinopathy of Prematurity: Implications for Vascular Regression, Invest. Ophthalmol. Vis. Sci. 45 (2004) 2049–2055. 10.1167/iovs.03-1346.

[20] L.C. Zepeda-Romero, M. Vazquez-Membrillo, E. Adan-Castro, F. Gomez-Aguayo, J.A. Gutierrez-Padilla, E. Angulo-Castellanos, J.C. Barrera de Leon, C. Gonzalez-Bernal, M.A. Quezada-Chalita, A. Meza-Anguiano, N. Diaz-Lezama, G. Martinez de la Escalera, J. Triebel, C. Clapp, Higher prolactin and vasoinhibin serum levels associated with incidence and progression of retinopathy of prematurity, Pediatr. Res. 81 (2017) 473–479. 10.1038/pr.2016.241.

[21] A. Hellström, L.E. Smith, O. Dammann, Retinopathy of prematurity, The Lancet 382 (2013) 1445–1457. 10.1016/S0140-6736(13)60178-6.

[22] F. Gross, J. Lazar, H. Orth, Inhibition of the Renin-Angiotensinogen Reaction by Pepstatin, Science 175 (1972) 656–656. 10.1126/science.175.4022.656.

[23] F. Suzuki, K. Murakami, Y. Nakamura, T. Inagami, 10 - Renin, in: A.J. Barrett, N.D. Rawlings, J.F. Woessner (Eds.), Handb. Proteolytic Enzym. Second Ed., Academic Press, London, 2004: pp. 54–61. 10.1016/B978-0-12-079611-3.50018-5.

[24] A.K. Kanugula, J. Kaur, J. Batra, A.R. Ankireddypalli, R. Velagapudi, Renin-Angiotensin System: Updated Understanding and Role in Physiological and Pathophysiological States, Cureus 15 (2023) e40725. 10.7759/cureus.40725.

[25] M. Holappa, H. Vapaatalo, A. Vaajanen, Many Faces of Renin-angiotensin System - Focus on Eye, Open Ophthalmol. J. 11 (2017) 122–142. 10.2174/1874364101711010122.

[26] M. Nath, P. Chandra, N. Halder, B. Singh, A.K. Deorari, A. Kumar, R. Azad, T. Velpandian, Involvement of Renin-Angiotensin System in Retinopathy of Prematurity - A Possible Target for Therapeutic Intervention, PLOS ONE 11 (2016) e0168809. 10.1371/journal.pone.0168809.

[27] C. Clapp, F.J. Lopez-Gomez, G. Nava, A. Corbacho, L. Torner, Y. Macotela, Z. Duenas, A. Ochoa, G. Noris, E. Acosta, E. Garay, G.M. de la Escalera, Expression of prolactin mRNA and of prolactin-like proteins in endothelial cells: evidence for autocrine effects, J. Endocrinol. 158 (1998) 137–144. 10.1677/joe.0.1580137.

[28] B. Staindl, P. Berger, R. Kofler, G. Wick, Monoclonal antibodies against human, bovine and rat prolactin: epitope mapping of human prolactin and development of a two-site immunoradiometric assay, J. Endocrinol. 114 (1987) 311–318. 10.1677/joe.0.1140311.

[29] P. Saftig, M. Hetman, W. Schmahl, K. Weber, L. Heine, H. Mossmann, A. Köster, B. Hess, M. Evers, K. von Figura, Mice deficient for the lysosomal proteinase cathepsin D exhibit progressive atrophy of the intestinal mucosa and profound destruction of lymphoid cells., EMBO J. 14 (1995) 3599–3608. 10.1002/j.1460-2075.1995.tb00029.x.

[30] A.M. Waterhouse, J.B. Procter, D.M.A. Martin, M. Clamp, G.J. Barton, Jalview Version 2—a multiple sequence alignment editor and analysis workbench, Bioinformatics 25 (2009) 1189–1191. 10.1093/bioinformatics/btp033.

[31] G.E. Crooks, G. Hon, J.-M. Chandonia, S.E. Brenner, WebLogo: A Sequence Logo Generator, Genome Res. 14 (2004) 1188–1190. 10.1101/gr.849004.

[32] R. Di Nicolantonio, F.A. Mendelsohn, Plasma renin and angiotensin in dehydrated and rehydrated rats, Am. J. Physiol.-Regul. Integr. Comp. Physiol. 250 (1986) R898–R901. 10.1152/ajpregu.1986.250.5.R898.

[33] E. Adán-Castro, L. Siqueiros-Márquez, G. Ramírez-Hernández, N. Díaz-Lezama, X. Ruíz-Herrera, F.F. Núñez, C.D. Núñez-Amaro, Ma.L. Robles-Osorio, T. Bertsch, J. Triebel, G. Martínez de la Escalera, C. Clapp, Sulpiride-induced hyperprolactinaemia increases retinal vasoinhibin and protects against diabetic retinopathy in rats, J. Neuroendocrinol. 34 (2022) e13091. 10.1111/jne.13091.

[34] A. Guillou, N. Romanò, F. Steyn, K. Abitbol, P. Le Tissier, X. Bonnefont, C. Chen, P. Mollard, A.O. Martin, Assessment of Lactotroph Axis Functionality in Mice: Longitudinal Monitoring of PRL Secretion by Ultrasensitive-ELISA, Endocrinology 156 (2015) 1924–1930. 10.1210/en.2014-1571.

[35] J.P. Robles, M. Zamora, L. Siqueiros-Marquez, E. Adan-Castro, G. Ramirez-Hernandez, F.F. Nuñez, F. Lopez-Casillas, R.P. Millar, T. Bertsch, G. Martínez de la Escalera, J. Triebel, C. Clapp, The HGR motif is the antiangiogenic determinant of vasoinhibin: implications for a therapeutic orally active oligopeptide, Angiogenesis 25 (2022) 57–70. 10.1007/s10456-021-09800-x.

[36] L. Jia, R.D. Simpson, J. Yuan, Z. Xu, W. Zhao, S. Cacatian, C.M. Tice, J. Guo, A. Ishchenko, S.B. Singh, Z. Wu, B.M. McKeever, Y. Bukhtiyarov, J.A. Johnson, C.P. Doe, R.K. Harrison, G.M. McGeehan, L.W. Dillard, J.J. Baldwin, D.A. Claremon, Discovery of VTP-27999, an Alkyl Amine Renin Inhibitor with Potential for Clinical Utility, ACS Med. Chem. Lett. 2 (2011) 747–751. 10.1021/ml200137x.

[37] T. Nakagawa, J. Akaki, R. Satou, M. Takaya, H. Iwata, A. Katsurada, K. Nishiuchi, Y. Ohmura, F. Suzuki, Y. Nakamura, The His-Pro-Phe motif of angiotensinogen is a crucial determinant of the substrate specificity of renin, 388 (2007) 237–246. 10.1515/BC.2007.026.

[38] A. Martyniak, P.J. Tomasik, A New Perspective on the Renin-Angiotensin System, Diagnostics 13 (2023) 16. 10.3390/diagnostics13010016.

[39] J. Triebel, T. Bertsch, C. Bollheimer, D. Rios-Barrera, C.F. Pearce, M. Hüfner, G. Martínez de la Escalera, C. Clapp, Principles of the prolactin/vasoinhibin axis, Am. J. Physiol.-Regul. Integr. Comp. Physiol. 309 (2015) R1193–R1203. 10.1152/ajpregu.00256.2015.

[40] J.L. Berka, A.J. Stubbs, D.Z. Wang, R. DiNicolantonio, D. Alcorn, D.J. Campbell, S.L. Skinner, Renin-containing Müller cells of the retina display endocrine features., Investigative Ophthalmology & Visual Science 36 (1995) 1450–1458.

[41] A.J. White, S.C. Cheruvu, M. Sarris, S.S. Liyanage, E. Lumbers, J. Chui, D. Wakefield, P.J. McCluskey, Expression of classical components of the renin-angiotensin system in the human eye, J Renin Angiotensin Aldosterone Syst 16 (2015) 59–66. 10.1177/1470320314549791.

[42] C. Clapp, L. Martínez de la Escalera, G. Martínez de la Escalera, Prolactin and blood vessels: A comparative endocrinology perspective, Gen. Comp. Endocrinol. 176 (2012) 336–340. 10.1016/j.ygcen.2011.12.033.

[43] H. Nishimura, M.L.S. Sequeira-Lopez, Phylogeny and ontogeny of the renin-angiotensin system: Current view and perspectives, Comp. Biochem. Physiol. A. Mol. Integr. Physiol. 254 (2021) 110879. 10.1016/j.cbpa.2020.110879.

[44] S. Marshall, M. Gelato, J. Meites, Serum Prolactin Levels and Prolactin Binding Activity in Adrenals and Kidneys of Male Rats After Dehydration, Salt Loading, and Unilateral Nephrectomy, Exp. Biol. Med. 149 (1975) 185–188. 10.3181/00379727-149-38769.

[45] S. Kaufman, B.J. Mackay, Plasma prolactin levels and body fluid deficits in the rat: causal interactions and control of water intake., J. Physiol. 336 (1983) 73–81. 10.1113/jphysiol.1983.sp014567.

[46] A.S. Chang, R. Grant, H. Tomita, H.-S. Kim, O. Smithies, M. Kakoki, Prolactin alters blood pressure by modulating the activity of endothelial nitric oxide synthase, Proc. Natl. Acad. Sci. 113 (2016) 12538–12543. 10.1073/pnas.1615051113.

[47] C.D. Nuñez-Amaro, A.I. Moreno-Vega, E. Adan-Castro, M. Zamora, R. Garcia-Franco, P. Ramirez-Neria, M. Garcia-Roa, Y. Villalpando, J.P. Robles, G. Ramirez-Hernandez, M. Lopez, J. Sanchez, E. Lopez-Star, T. Bertsch, G. Martinez de la Escalera, Ma.L. Robles-Osorio, J. Triebel, C. Clapp, Levosulpiride Increases the Levels of Prolactin and Antiangiogenic Vasoinhibin in the Vitreous of Patients with Proliferative Diabetic Retinopathy, Transl. Vis. Sci. Technol. 9 (2020) 27. 10.1167/tvst.9.9.27.

[48] M.S. Kuchay, A. Mithal, Levosulpiride and Serum Prolactin Levels, Indian J. Endocrinol. Metab. 21 (2017) 355. 10.4103/ijem.IJEM_555_16.

[49] C. Molinari, E. Grossini, D.A.S.G. Mary, F. Uberti, E. Ghigo, F. Ribichini, N. Surico, G. Vacca, Prolactin Induces Regional Vasoconstriction through the β2-Adrenergic and Nitric Oxide Mechanisms, Endocrinology 148 (2007) 4080–4090. 10.1210/en.2006-1577.

[50] C. Gonzalez, A.M. Corbacho, J.P. Eiserich, C. Garcia, F. Lopez-Barrera, V. Morales-Tlalpan, A. Barajas-Espinosa, M. Diaz-Muñoz, R. Rubio, S.-H. Lin, G. Martinez de la Escalera, C. Clapp, 16K-Prolactin Inhibits Activation of Endothelial Nitric Oxide Synthase, Intracellular Calcium Mobilization, and Endothelium-Dependent Vasorelaxation, Endocrinology 145 (2004) 5714–5722. 10.1210/en.2004-0647.

[51] A.D. de Kloet, S. Pitra, L. Wang, H. Hiller, D.J. Pioquinto, J.A. Smith, C. Sumners, J.E. Stern, E.G. Krause, Angiotensin Type-2 Receptors Influence the Activity of Vasopressin Neurons in the Paraventricular Nucleus of the Hypothalamus in Male Mice, Endocrinology 157 (2016) 3167–3180. 10.1210/en.2016-1131.

[52] S. Mejía, L.M. Torner, M.C. Jeziorski, C. Gonzalez, M.A. Morales, G.M. de la Escalera, C. Clapp, Prolactin and 16K prolactin stimulate release of vasopressin by a direct effect on hypothalamo-neurohypophyseal system, Endocrine 20 (2003) 155–161. 10.1385/ENDO:20:1-2:155.

[53] S.P. Tabruyn, C. Sabatel, N.-Q.-N. Nguyen, C. Verhaeghe, K. Castermans, L. Malvaux, A.W. Griffioen, J.A. Martial, I. Struman, The Angiostatic 16K Human Prolactin Overcomes Endothelial Cell Anergy and Promotes Leukocyte Infiltration via Nuclear Factor-κB Activation, Mol. Endocrinol. 21 (2007) 1422–1429. 10.1210/me.2007-0021.

[54] G. Ortiz, M.G. Ledesma-Colunga, Z. Wu, J.F. García-Rodrigo, N. Adan, O.F. Martinez-Diaz, E.A. De Los Ríos, F. López-Barrera, G. Martínez de la Escalera, C. Clapp, Vasoinhibin is Generated and Promotes Inflammation in Mild Antigen-induced Arthritis, Endocrinology 163 (2022) bqac036. 10.1210/endocr/bqac036.

[55] C.J. Moravski, D.J. Kelly, M.E. Cooper, R.E. Gilbert, J.F. Bertram, S. Shahinfar, S.L. Skinner, J.L. Wilkinson-Berka, Retinal Neovascularization Is Prevented by Blockade of the Renin-Angiotensin System, Hypertension 36 (2000) 1099–1104. 10.1161/01.HYP.36.6.1099.

